# Kappa opioid receptors regulate cocaine effects on nucleus accumbens dopamine through phosphorylation of dopamine transporter at the threonine 53 site

**DOI:** 10.64898/2026.05.06.722744

**Authors:** Emanuel F Lopes, Paige M Estave, Alyson M Curry, Kathryn R Beard, Monica H Dawes, Jonathan H Sciortino, Kathleen A Grant, Lankupalle D Jayanthi, Sammanda Ramamoorthy, Katherine M Holleran, Sara R Jones

**Author notes:** **Correspondence:** Sara R Jones, Department of Translational Neuroscience, WFUSM, Medical Center Blvd., Winston-Salem, NC 27157.

## Abstract

Chronic cocaine exposure leads to tolerance of the dopamine transporter (DAT) to cocaine and heightened kappa opioid receptor (KOR)-induced inhibition of dopamine (DA) release in the nucleus accumbens (NAc). KORs on presynaptic DA terminals inhibit DA release and modulate DAT function, but whether KOR signaling causes DAT tolerance is unknown. We demonstrate that rats self-administering cocaine escalate their intake, and the KOR antagonist norbinaltorphimine (nBNI) blunts this escalation. We employed fast-scan cyclic voltammetry (FSCV) in acute brain slices containing the NAc core from these rats and found that the cocaine exposure paradigm resulted in tolerance of the DAT to cocaine, an effect absent in animals pre-treated with nBNI. In drug-naïve mice, either superfusion of the KOR agonist U50,488 or a single *in vivo* injection of U69,593 inhibited DA release and reduced cocaine-induced inhibition of DA reuptake, reproducing the tolerance phenotype without any drug history. We replicated this finding in the NAc of rats, and with the uptake inhibitor nomifensine in rhesus macaques, indicating conservation across rodents and non-human primates. As KOR activation results in phosphorylation of the threonine-53 site on the DAT, we tested whether this is required for the KOR-mediated DAT tolerance to cocaine. We show that a mouse line with a phosphorylation-insensitive alanine-53 mutation of the DAT has enhanced DA release and uptake. Remarkably, KOR agonism no longer mediated DAT tolerance to cocaine in these mice, demonstrating the dependence of this effect on the threonine-53 site and highlighting a KOR-driven mechanism underlying cocaine tolerance at the DAT.

## Introduction

Cocaine misuse is an escalating public health crisis with profound and far-reaching effects on individuals and society ^1^. Cocaine use heightens the risk of overdose, cardiovascular and respiratory disease, and neuropsychiatric conditions ^2,3^. Repeated cocaine use can escalate into cocaine use disorder (CUD), defined as the inability to control cocaine use despite adverse consequences. To date, the FDA has not approved any medication to treat CUD. Therefore, a greater understanding of cocaine’s mechanisms and the neurobiological underpinnings of CUD are crucial to facilitate the development of effective therapeutic interventions.

Cocaine primarily exerts its reinforcing effects by increasing dopamine (DA) signaling in the nucleus accumbens (NAc) by inhibiting DA reuptake through the dopamine transporter (DAT) ^4–8^. Chronic cocaine use leads to a significant dysregulation of DA signaling. Of particular interest to this study, clinical and preclinical studies have found that cocaine-induced DA uptake inhibition is blunted ^9–13^, i.e., there is a tolerance of the DAT to cocaine. This neurobiological tolerance has been linked to the development of tolerance to behavioral effects, leading to increases in response behavior as progressively larger quantities of the drug are required to obtain and maintain the desired drug effects ^14–17^.

Our lab has further shown that cocaine exposure heightens kappa opioid receptor (KOR) agonist-induced inhibition of DA release in the NAc ^18^. The dynorphin/KOR (Dyn/KOR) system is a critical modulator of stress, pain, and reward ^19,20^. While expressed widely in the central nervous system, KORs are particularly enriched in the ventral tegmental area (VTA) and the NAc, the principal regions of the dopaminergic mesolimbic pathway ^21,22^. The Gi-linked KOR is thought to mediate a net inhibitory influence on DA levels in the NAc by (1) inhibiting terminal DA release ^23–29^, (2) inhibiting VTA DA neuron firing ^26,30,31^, and (3) increasing DA uptake rates ^26,32–34^, though this effect may depend on experimental method and treatment regimen ^35,36^. Dyn levels and/or KOR activity are increased in the mesolimbic DA pathway following stress ^37–39^, pain ^40,41^, and exposure to drugs of abuse such as cocaine ^10,18^, amphetamine ^42,43^, ethanol ^24,27,44,45^, and opioids ^40,46^. Consistent with the inhibitory actions of KORs on DA release, stress and pain are associated with mesolimbic hypodopaminergia and increases in function or expression of the Dyn/KOR system have been found to drive negative affect ^47,48^.

The dynamics of vesicular DA release and its clearance determine the availability of extracellular DA for dopaminergic transmission. KOR is expressed in DA terminals along with DAT ^49^, and the ability of the KOR to modify DA uptake lies in its indirect interactions with the DAT, a primary regulator of extracellular DA levels ^50^. DAT function is tightly regulated by its phosphorylation state, which is controlled by intracellular signaling pathways including protein kinase C and extracellular signal-regulated kinase (ERK; reviewed in ^51–55^). Indeed, KOR activation has been shown to result in the phosphorylation of the threonine-53 residue of the DAT (pT53-DAT) involving ERK1/2 ^32,34,56,57^. pT53 is involved in modulating DAT activity and surface expression ^34^, and mice lacking this DAT phosphorylation site (DAT-T53A mice) exhibit reduced sensitivity to cocaine-mediated hyperlocomotion and synaptosomal DA uptake inhibition ^57^. DAT-T53A mice are also insensitive to KOR-mediated increases in synaptosomal DA uptake, DAT surface expression, locomotor suppression, conditioned place aversion, and KOR-mediated enhanced cocaine preference ^56^. Consequently, pT53-DAT is a crucial point of interaction between Dyn/KOR signaling and functional regulations of the DAT and is central to many KOR and cocaine-mediated behaviors.

In this study, we examined whether acute activation of the KOR is sufficient to modify cocaine-mediated inhibition of DA uptake and further tested whether this effect requires pT53-DAT. We demonstrated that in rats self-administering cocaine in an extended access, limited intake procedure, the KOR antagonist nBNI blunts escalation of cocaine intake. We then employed fast-scan cyclic voltammetry (FSCV) in acute brain slices containing the NAc core and found that the cocaine exposure paradigm did not limit the effect of cocaine on the DAT in animals pre-treated with nBNI. We then showed in drug-naïve mice that KOR activation *in vivo* and in acute striatal slices can suppress cocaine-mediated inhibition of DA uptake, and that this effect can be replicated in non-human primate (rhesus macaque) NAc with another DAT inhibitor, nomifensine. We then tested these mechanisms on DAT-T53A mice. DAT-T53A mice showed higher evoked DA release and uptake and reduced KOR-effects on peak evoked DA. Remarkably, in DAT-T53A mice, KOR activation no longer induced DAT tolerance to cocaine, demonstrating that this effect requires pT53-DAT and suggesting a KOR-driven molecular pathway that underlies cocaine tolerance at the DAT.

## Materials and Methods

### Animals

Male Sprague-Dawley rats (325–400 g; Envigo, Indianapolis, IN, USA) were maintained on a 12:12 h light/dark cycle (lights on: 1500) and given food and water *ad libitum.* Wake Forest University School of Medicine’s (WFUSM) Institutional Animal Care and Use Committee (IACUC) approved the experimental protocol. Animals were maintained according to the National Institutes of Health guidelines in Association for Assessment and Accreditation of Laboratory Animal Care accredited facilities.

Male and female homozygous DAT-T53A and WT C57BL/6J mice (8-10 weeks old) were bred in-house for this study. DAT-T53A mice were generated on a C57BL/6J background through a CRISPR/Cas9 approach and are viable with no significant abnormalities in male or female mice regarding growth, bodyweight, body temperature, or general health ^57^. Mice were group housed, on a 12:12 h light cycle (lights on: 0600), with all experiments performed in the light cycle with food *ad libitum*. The protocol was approved by WFUSM’s IACUC. All methods were performed in accordance with the relevant guidelines and regulations.

Young adult male and female rhesus macaques (N = 3 male and 4 female) were housed in quadrant cages (0.8×0.8×0.9 m) and maintained on an 11-hour light cycle (ON: 0800) at constant temperature (20-22 °C) and humidity (65%). All experimental protocols were carried out in accordance with the Guide for the Care and Use of Laboratory Animals and approved by the Oregon National Primate Research Center IACUC. Rhesus monkeys used in this study were drug- and alcohol-naïve, and received training for awake blood draws, medical check-ups, and MRI imaging ^58^.

### Catheter implantation

Rats were implanted with a chronic indwelling jugular catheter under ketamine (100 mg/kg; i.p.) and xylazine (8 mg/kg; i.p.) anasthesia, with ketoprofen (5 mg/kg; s.c.) for analgesia, as described previously ^59^. After surgery, animals were single housed in custom-made operant conditioning chambers (30×30×30 cm), which served as both a housing and experimental chamber. Self-administration sessions occurred in the active/dark cycle (0900-1500). All control animals were implanted with a closed-off port, which allowed animals to be tethered in their cage to control for surgery and housing conditions.

### Extended Access, Limited Intake Procedure

Prior to cocaine self-administration, rats were randomly assigned to receive an injection of nBNI (10 mg/kg; i.p.) or vehicle (sterile injectable water) 18 hours prior to their first cocaine self-administration session. Two days following surgery, animals self-administered cocaine (1.5 mg/kg/inf; ∼4 sec/inf) on a fixed-ratio one (FR1) schedule of reinforcement. Animals had no prior operant training. Each response on the lever resulted in a cocaine infusion, retraction of the lever and illumination of a stimulus light during the infusion. Sessions were terminated after 40 infusions were reached or 6 hours elapsed, whichever occurred first. Animals were required to complete 40 infusions for 5 consecutive days. nBNI or vehicle was administered every 7 days until animals completed this requirement.

### *Ex vivo* fast scan cyclic voltammetry (FSCV)

*Ex vivo* FSCV was used to assess alterations in DA dynamics in the NAc core as previously reported ^60^. Briefly, mice or rats were rapidly decapitated and brains extracted. Coronal brain slices containing the NAc core (300 µm, mice; 400 µm, rats) were prepared using a vibrating tissue slicer and incubated in oxygenated artificial cerebrospinal fluid (aCSF) at room temperature that contained (in mM): NaCl (126), KCl (2.5), NaH_2_PO_4_ (1.2), CaCl_2_ (2.4), MgCl_2_ (1.2), NaHCO_3_ (25), glucose (11), L-ascorbic acid (0.4). Once slices were transferred to recording chambers, a carbon fiber microelectrode (100–200 μm length, 7 μm diameter) and a bipolar stimulating electrode were placed into the NAc core. DA release was evoked every 3 min by applying a single electrical pulse from the bipolar stimulating electrode (750 μA, 4 ms, monophasic) to the tissue. The DA concentration was recorded by applying a triangular waveform (−0.4 to +1.2 to −0.4 V vs. Ag/AgCl, at a rate of 400 V/s) to the carbon fiber microelectrode. Once the evoked release of DA was stable, drugs were diluted into the superfusion buffer and release events used for subsequent analysis were selected once the effect of each drug reached stability (∼45 min and <10% peak height variability across 3 consecutive collections).

For mouse pre-treatment experiments, on each experimental day a pair of mice was injected with either U69,593 (s.c., 0.1 mg/kg), or vehicle (s.c., 20% propylene glycol in saline). One hour after injection, animals were rapidly decapitated and brains extracted as above.

For non-human primate FSCV experiments, animals were sedated with ketamine (10 mg/kg) and maintained in deep anesthesia via isoflurane for transcardial perfusion. Following completion of the perfusion, brains were removed and placed into a brain block for dissection into 4 mm coronal sections, as described previously ^61^. 250 µm thick coronal brain slices containing the NAc were incubated in oxygenated aCSF at room temperature. A carbon fiber microelectrode (100-200 µm length, 7 µm diameter) and bipolar stimulating electrode were placed in close proximity on the tissue. DA release and detection were performed as described in mouse FSCV experiments. DA release experiments were conducted following establishment of a stable baseline.

To evaluate DA release and uptake kinetics, the program Demon Voltammetry and Analysis was used ^62^. Individual electrode calibration factors were calculated as described previously ^63^. Michaelis–Menten based modeling in Demon Voltammetry and Analysis was then used to determine the amount of stimulated DA release, the maximal rate of DA uptake (V_max_), and the ability of DA to bind to the DAT in the presence of a competitive inhibitor such as cocaine (apparent K_m_).

### Drugs

Cocaine ((−)-cocaine HCl, RTI Log No.: 14201-134), U50,488H (U50,488; RTI Log No.: 13692-54B), and nBNI (norbinaltorphimine dihydrochloride; RTI Log No.: 14055-160) were generously provided by the NIDA Drug Supply Program and dissolved in deionized water for FSCV experiments. Nomifensine maleate was purchased from Tocris Bioscience (Minneapolis, MN, USA) and dissolved in dimethyl sulfoxide. For self-administration experiments, cocaine was dissolved in sterile saline and nBNI was dissolved in sterile injectable water.

### Experimental design and statistical analysis

GraphPad Prism 11 (Dotmatics/GraphPad, Boston, MA) was used to conduct all analyses. Data are reported as mean ± standard error, with “n” referring to the number of slices and “N” referring to the number of animals. The significance level was set at p < 0.05. Comparisons for statistical significance were assessed by one- and two-way ANOVA with Šídák’s or Tukey’s post hoc tests, unpaired t-tests, and paired t-tests where appropriate. In datasets with male and female animals, sex effects were tested using 2-way ANOVAs for peak [DA] or V_max_, and 3-way ANOVAs for K_m_ changes in the presence of uptake inhibitors. Where not reported, no significant effects were detected.

## Results

### Treatment with the KOR antagonist nBNI blunts escalation of cocaine intake rate in rats and prevents changes to the effects of cocaine on the DAT

We have previously shown that 5 days of 40 cocaine infusions (1.5 mg/kg/inf) results in an escalated rate of intake across cocaine self-administration (SA) sessions ^11,64,65^. This escalation can be expressed as a decrease in the average interinfusion interval between the 40 infusions. We aimed to assess the role of KORs in this escalation effect by pre-treating rats with nBNI (10 mg/kg, i.p.) or vehicle prior to their initial cocaine self-administration session (**Figure 1A**). There was a significant effect of session, indicating animals escalate cocaine consumption (**Figure 1B**, repeated measures two-way ANOVA, effect of session: *F*(4,79) = 24.44, *p* < 0.0001, *N* = 12 vehicle, 12 nBNI rats), and nBNI significantly decreased intake rate (**Figure 1B**, repeated measures two-way ANOVA, effect of treatment: *F*(1,20) = 5.990, *p* = 0.0237), with a significant increase in interinfusion interval on day 4 between groups (**Figure 1B**, Šídák’s multiple comparisons test, *p* = 0.0284). When comparing the average interinfusion interval between groups across the 5 sessions, nBNI-treated animals show a significant increase in interinfusion interval (**Figure 1C**, unpaired t-test: *t*(22) = 3.073, *p* = 0.0056). Consequently, blocking KORs is sufficient to blunt escalation of cocaine intake rate during cocaine SA sessions.

**Figure 1.**
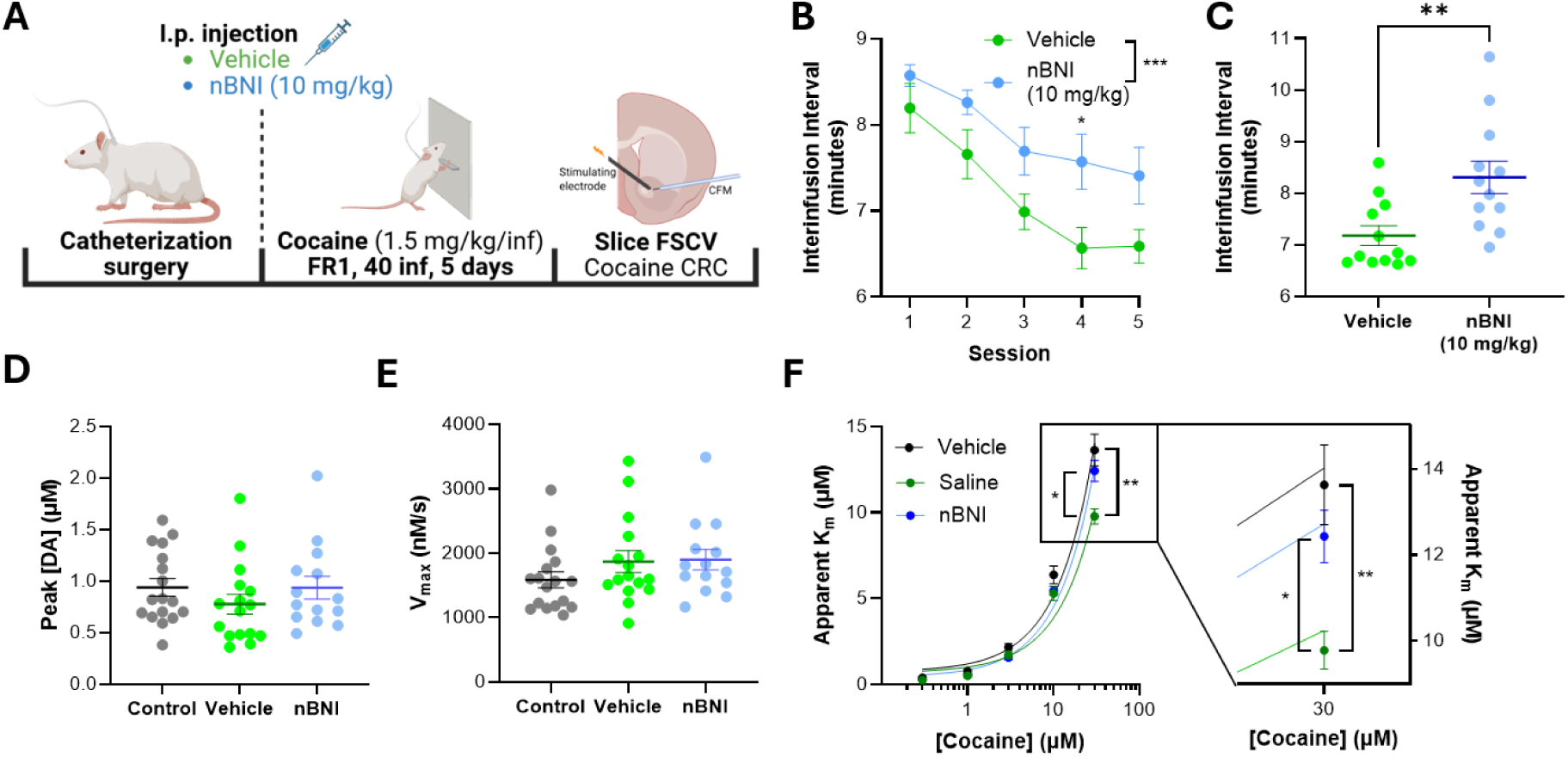
- Treatment with a KOR antagonist blocks escalation of cocaine intake rate and cocaine tolerance at the DAT. **(A)** Schematic of experimental timeline. **(B)** Average ± SEM cocaine interinfusion interval (time between each infusion) throughout the 5 cocaine self-administration sessions in animals treated with vehicle (green) or nBNI (blue). **(C)** Average cocaine interinfusion interval per animal, collapsed across all sessions. N = 12 vehicle + 12 nBNI pre-treated rats. **(D)** Average ± SEM peak evoked [DA] in acute rat NAc slices following cocaine self-administration in sham control rats (black), vehicle pre-treated rats (green), and rats pre-treated with nBNI (blue - 10 mg/kg). **(E)** Maximal velocity of DA uptake (V_max_) in control, vehicle-pretreated, and nBNI-pretreated rats following cocaine self-administration. **(F)** Left: Average apparent K_m_ across increasing concentrations of cocaine (0.3, 1, 3, 10, 30 μM). Right: Focused view of data at 30 μM cocaine. n = 15 (control), 16 (vehicle), 16 (nBNI) slices, N = 11 (control), 9 (vehicle), 12 (nBNI) rats. * p ≤ 0.05, ** p ≤ 0.01, *** p ≤ 0.001, **** p ≤ 0.0001.

Our laboratory has shown that rats self-administering cocaine on an extended-access, limited-intake procedure show a reduced effect of cocaine on the DAT ^9,11,64,66^. We will subsequently refer to this reduced effect of cocaine on the DAT as DAT “tolerance” to cocaine. As KOR inhibition reduced cocaine intake rate, we then examined whether nBNI also alters the emergence of DAT tolerance to cocaine. To address these questions and probe NAc core DA dynamics, we employed FSCV in acute rat NAc slices to detect extracellular DA concentration ([DA]) evoked by a single electrical pulse. We sampled [DA] release from sham surgery animals not exposed to cocaine (control), from animals exposed to the cocaine self-administration procedure pre-treated with vehicle, and animals exposed to the cocaine self-administration paradigm pre-treated with nBNI. Peak evoked [DA] (**Figure 1D**, one-way ANOVA: *F*(2,44) = 0.935, *p* = 0.40) and uptake (V_max_, **Figure 1E**, one-way ANOVA: *F* (2,44) = 1.365, *p* = 0.27) were not significantly different between the three treatment conditions. We further superfused increasing concentrations of cocaine on acute slices from animals exposed to the three conditions and assessed the ability of cocaine to inhibit DA uptake (**Figure 1F**, apparent K_m_ ^62^). We found a significant effect of treatment (**Figure 1F**, mixed-effects analysis, effect of treatment: *F* (1.778,28.45) = 6.702, *p* = 0.0054, *n* = 15 control, 16 vehicle, 16 nBNI slices) with significant post hoc differences at 30 μM cocaine between control and vehicle conditions (**Figure 1F**, Tukey’s multiple comparisons test: *p* = 0.0138), and nBNI and vehicle conditions (**Figure 1F**, Tukey’s multiple comparisons test: *p* = 0.0036). This suggests that blocking KORs prevents the impact of the cocaine exposure paradigm in causing DAT tolerance. Put together, these data suggest that inhibiting KORs prevents escalation of cocaine use, and the development of DAT tolerance to cocaine.

### *In vivo* pretreatment with KOR agonist induces DAT tolerance in drug-naïve mice

We demonstrated that repeated cocaine self-administration resulted in DAT tolerance to cocaine, an effect blocked by treatment with nBNI. We then tested whether a single *in vivo* dose of a KOR agonist is sufficient to mediate DAT tolerance in drug-naïve animals. We treated a cohort of male C57BL/6J mice *in vivo* with the KOR agonist U69,593 (s.c., 0.1 mg/kg, **Figure 2A**). Following a single subcutaneous injection of vehicle or U69,593 and a 1-hour incubation period, we obtained acute striatal slices from these animals. *In vivo* pre-treatment with U69,593 suppressed peak [DA] (**Figure 2B**, unpaired t-test: *t*(22) = 2.227, *p* = 0.0365, *n* = 13 vehicle and 11 U69,593 slices). Uptake was not significantly different following U69,593 pretreatment (**Figure 2C**, WT: unpaired t-test: *t*(22) = 1.152, *p* = 0.2618, *n* = 13 vehicle and 11 U69,593 slices). Finally, the effects of cocaine were reduced in WT animals following U69,593 pretreatment compared to vehicle-treated animals (**Figure 2D**, two-way ANOVA, effect of U69,593: *F*(1,32) = 8.184, *p* = 0.0074, *n* = 5 vehicle and 5 U69,593 slices). To test whether any residual U69,593 may be activating KORs on the acute slices, we superfused the KOR antagonist nBNI (10 µM) in a subset of slices. We found no detectable nBNI-sensitive component to peak [DA] between slices from vehicle- or U69,593-pretreated mice (**Figure 2E**, unpaired t-test, *t*(10) = 0.6330, *p* = 0.5410). Together, these data show a single *in vivo* dose of a KOR agonist is sufficient to mediate DAT tolerance to cocaine.

**Figure 2.**
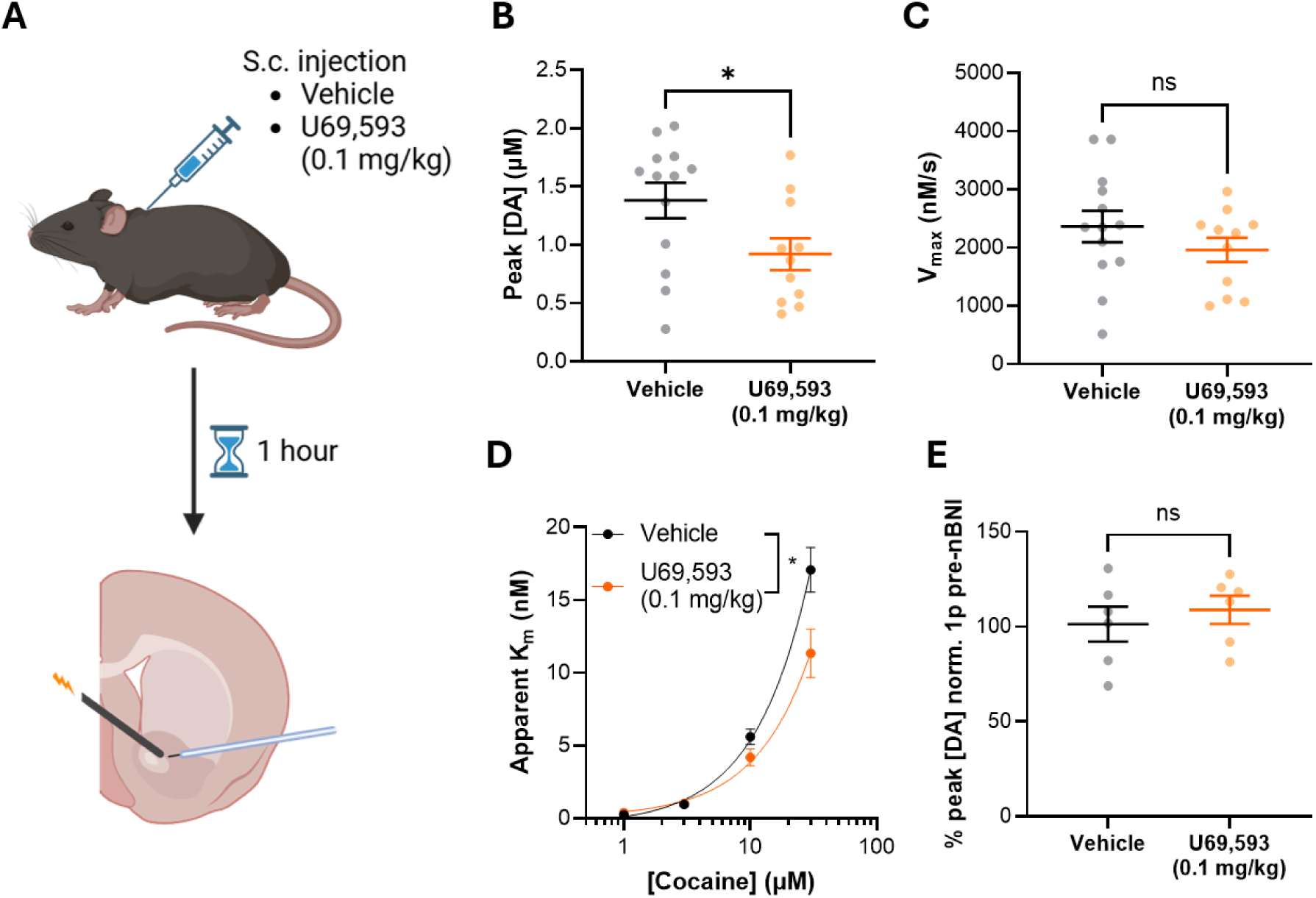
- *In vivo* pretreatment with KOR agonist induces DAT tolerance to cocaine in drug-naïve WT mice. **(A)** Schematic of experimental set-up. Male WT mice were injected with the KOR agonist U69,593 (0.1 mg/kg) or vehicle (20% propylene glycol in saline). Following a 1 hour period, animals were sacrificed and acute NAc core striatal slices were obtained for FSCV. **(B)** Average ± SEM peak evoked [DA] in vehicle (black) and U69,593 (yellow) conditions in acute slices from WT animals. **(C)** Maximal velocity of DA uptake (V_max_) in vehicle and U69,593 conditions. n = 13 vehicle and 11 U69,593 slices. **(D)** Average apparent K_m_ across cocaine concentrations in vehicle and U69,593 treated WT animals. n = 5 vehicle and 5 U69,593 slices. **(E)** Peak [DA] after 10 µM nBNI, expressed as a percentage of pre-nBNI baseline. n = 6 vehicle and 6 U69,593 slices, N = 5 vehicle and 5 U69,593 mice. * p ≤ 0.05.

### KOR activation in acute NAc slices suppresses dopamine release and induces DAT tolerance to cocaine in drug-naïve mice

We demonstrated that KOR activation *in vivo* is sufficient to mediate DAT tolerance to cocaine. To test whether NAc KORs are sufficient, as opposed to circuits elsewhere in the intact brain, we activated KORs in deafferented slices containing the NAc. We subsequently assessed the effects of an acute superfusion of the KOR agonist U50,488 (100 nM) on DA release in NAc slices from drug-naïve C57BL/6J mice (**Figure 3A**). As described previously ^23,24,44^, U50,488 suppressed peak [DA] (**Figure 3A,B**, paired t-test: *t*(20) = 7.184, *p* < 0.0001, *n* = 21) without significantly impacting the rate of DA uptake (**Figure 3A,C**, paired t-test: *t*(20) = 1.179, *p* = 0.2521).

**Figure 3.**
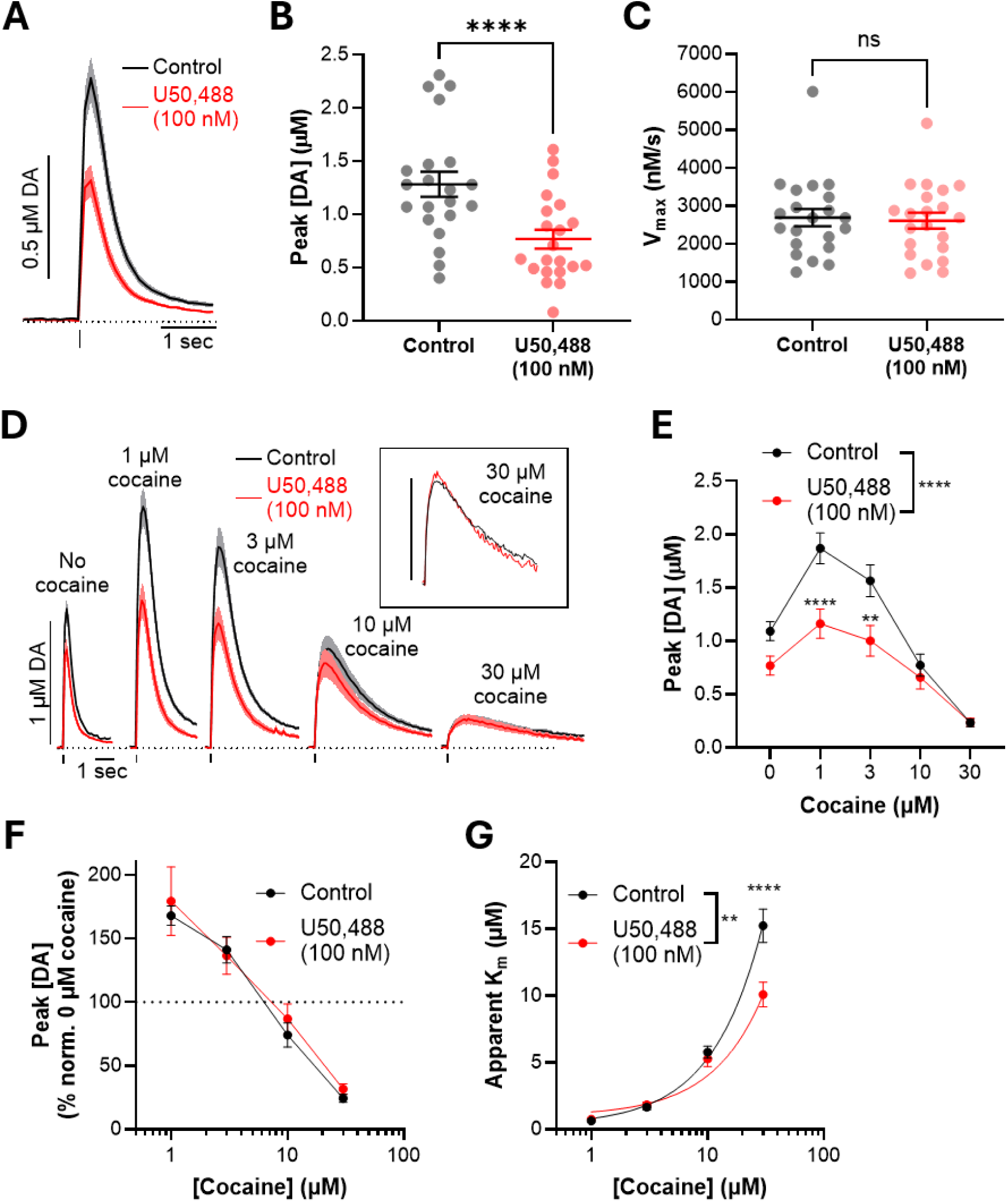
- KOR activation suppresses NAc core DA release and limits effects of cocaine. **(A)** Average ± SEM DA release events in drug-free conditions (Control, black line) and in the presence of the KOR agonist U50,488 (100 nM, red line). Vertical line indicates electrical stimulation timepoint, horizontal dotted line indicates 0 µM [DA]. **(B)** Average ± SEM peak evoked [DA] in drug-free and U50,488 conditions. **(C)** Maximal velocity of DA uptake (V_max_) in Control and U50,488 conditions. **(D)** Averaged DA release events evoked in increasing concentrations of cocaine (0, 1, 3, 10, 30 μM), in the presence or absence of U50,488. Inset: magnified representation of average DA release events in 30 μM cocaine. Bar = 0.2 μM [DA]. **(E)** Peak evoked [DA] across cocaine concentrations. **(F)** Peak evoked [DA] normalized to peak evoked [DA] at 0 μM cocaine across cocaine concentrations. **(G)** Average apparent K_m_ across cocaine concentrations. n = 21 slices, N = 6 male and 5 female mice. ** p ≤ 0.01, **** p ≤ 0.0001.

We subsequently superfused increasing concentrations of cocaine in the presence or absence of 100 nM U50,488 (**Figure 3D**). We then assessed changes to peak [DA] (**Figure 3E,F**) and the ability of cocaine to inhibit DA uptake (**Figure 3G**, apparent K_m_ ^62^). We found that U50,488 suppressed peak [DA] (**Figure 3E**, two-way ANOVA, effect of U50,488: *F*(1,200) = 23.38, *p* < 0.0001, *n* = 21), with significant post hoc effects at 1 μM and 3 μM cocaine (**Figure 3E**, Šídák’s multiple comparisons test, 1 μM cocaine: *p* < 0.0001, 3 μM cocaine: *p* = 0.0022). To test whether KOR agonism modified how cocaine impacts peak evoked [DA], we normalized peak [DA] values across the various cocaine concentrations to the peak [DA] value in the absence of cocaine, and found that U50,488 did not significantly alter how cocaine changes peak [DA] (**Figure 3F**, two-way ANOVA, effect of U50,488: *F*(1,160) = 0.6484, *p* = 0.4219, *n* = 21 control and U50,488 slices). When testing the ability of DA to bind to the DAT in the presence of cocaine, U50,488 significantly reduced apparent K_m_ (**Figure 3G**, two-way ANOVA, effect of U50,488: *F*(1,160) = 9.443, *p* = 0.0025, *n* = 21). This effect was particularly prominent at the highest concentration of cocaine (**Figure 3G**, Šídák’s multiple comparisons test, 30 μM cocaine: *p* < 0.0001). These findings suggest that KOR activation in acute striatal slices induces DAT tolerance to cocaine, indicating a local terminal mechanism that mediates this effect.

### KOR-mediated blunting of dopamine uptake inhibitor sensitivity is conserved in rodents and non-human primates

We then aimed to test whether activating KORs similarly induced DAT tolerance to cocaine in drug-naïve rats (**Figure 4A**). Superfusing U50,488 (300 nM) significantly suppressed peak evoked [DA] (**Figure 4B**, paired t-test: *t*(7) = 3.357, *p* = 0.0121, *n* = 8 slices). Superfusing increasing concentrations of cocaine in the presence or absence of 300 nM U50,488 revealed that U50,488 significantly reduced cocaine-mediated changes in apparent K_m_ (**Figure 4C**, two-way ANOVA: effect of U50,488: *F*(1,70) = 4.765, *p* = 0.0324, *n* = 8), with significant post hoc differences at 30 μM cocaine (**Figure 4C**, Šídák’s multiple comparisons test: *p* < 0.0001).

**Figure 4.**
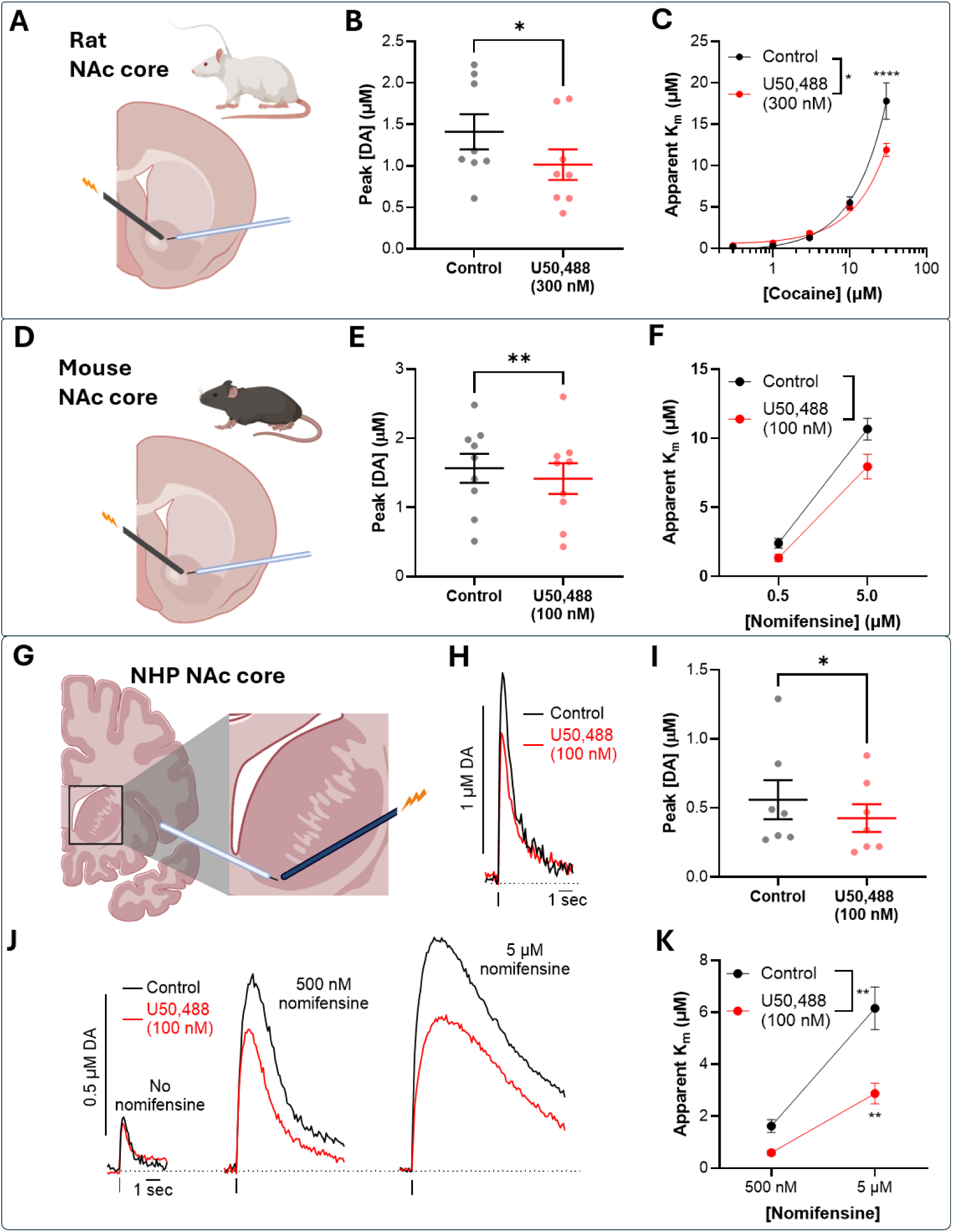
- KOR activation limits effects of dopamine reuptake inhibitors in rodents and NHPs. **(A)** Schematic describing FSCV experimental set up with acute rat NAc slices. **(B)** Peak evoked [DA] in drug-free (Ctrl, black) and U50,488 (red) conditions. **(C)** Average apparent K_m_ across cocaine concentrations (0.3, 1, 3, 10, 30 μM) in the presence (red) or absence (black) of 300 nM U50,488. n = 8 slices, N = 8 male rats. **(D)** Schematic describing FSCV experimental set up with acute mice slices. **(E)** Peak evoked [DA] in drug-free (Ctrl, black) and U50,488 (red) conditions. n = 9 slices. **(F)** Average apparent K_m_ across nomifensine concentrations (0.5 and 5 μM) in the presence (red) or absence (black) of 100 nM U50,488. n = 8 control and 9 U50,488 slices, N = 4 male and 4 female WT mice. **(G)** Acute NHP brain slices containing the NAc were obtained, and dopamine release was electrically evoked in the NAc with a bipolar stimulating electrode. DA release was detected through fast-scan cyclic voltammetry with a carbon-fiber microelectrode. **(H)** Representative DA release events, evoked by a single electrical pulse in drug-free conditions (Control, black line) and in the presence of the KOR agonist U50,488 (100 nM, red line). **(I)** Peak evoked [DA] in drug-free (Ctrl, black) and U50,488 (red) conditions. **(J)** Representative DA release events evoked in increasing concentrations of nomifensine (0, 0.5, 5 μM), in the presence (red) or absence (black) of 100 nM U50,488. **(K)** Average apparent K_m_ across nomifensine concentrations in the presence (red) or absence (black) of 100 nM U50,488. n = 7 slices, N = 3 male and 4 female rhesus monkeys. * p ≤ 0.05, ** p ≤ 0.01, *** p ≤ 0.001, *** p ≤ 0.0001.

To test whether the previously described mechanisms are evolutionarily conserved, we tested whether U50,488 can modify the effects of the dual DAT/norepinephrine transporter (DAT/NET) uptake inhibitor nomifensine on DA release in the NAc of rhesus macaques (referred to as non-human primates, NHP). Similar to cocaine, the ability of nomifensine to inhibit DA uptake is reduced in rats with a history of cocaine self-administration ^9^. To start, we questioned whether KOR activation limited the effects of nomifensine in the mouse NAc core (**Figure 4D-F**). U50,488 suppressed peak [DA] (**Figure 4E**, paired t-test: *t*(8) = 3.385, *p* = 0.0096, *n* = 9) and significantly limited the change in apparent K_m_ (**Figure 4F**, two-way ANOVA, effect of U50,488: *F*(1, 30) = 8.750, *p* = 0.0060, *n* = 8 control and 9 U50,488).

We then tested the effects of U50,488 on NHP NAc DA release (**Figure 4G**). U50,488 (100 nM) significantly suppressed peak [DA] (**Figure 4H,I**, paired t-test: *t*(6) = 2.713, *p* = 0.035, *n* = 7), mirroring the results seen in mouse NAc. Likewise, when sequentially superfusing nomifensine (500 nM and 5 μM), pre-incubation with U50,488 significantly limits the change in apparent K_m_ (**Figure 4J,K**, two-way ANOVA, effect of U50,488: *F*(1,6) = 14.07, *p* = 0.0095, *n* = 7). While no significant sex effects were detected in this dataset (**Figure 4K**, mixed-effects analysis, effect of sex: *F*(1,10) = 3.379, *p* = 0.0969), there was a significant interaction between sex and nomifensine (**Figure 4K**, mixed-effects analysis, sex x nomifensine: *F*(1,10) = 5.615, *p* = 0.0393). These findings indicate that KOR-dependent modulation of DAT tolerance is present in rodents and a primate species, supporting the translational relevance of this mechanism.

### Mice with phosphorylation-insensitive T53 residue of the DAT show alterations to baseline DA dynamics

We then explored mechanisms that may underlie the KOR-mediated inhibition of cocaine effects. T53 phosphorylation on the DAT is a major point of interaction between the KOR and the DAT and has been shown to regulate a number of KOR- and cocaine-related behaviors ^34,56,57^. We employed a mouse line with a substituted alanine at position 53 (DAT-T53A ^57^) to investigate the role of this phosphorylation site on the reduction of cocaine potency (**Figure 5A**).

**Figure 5.**
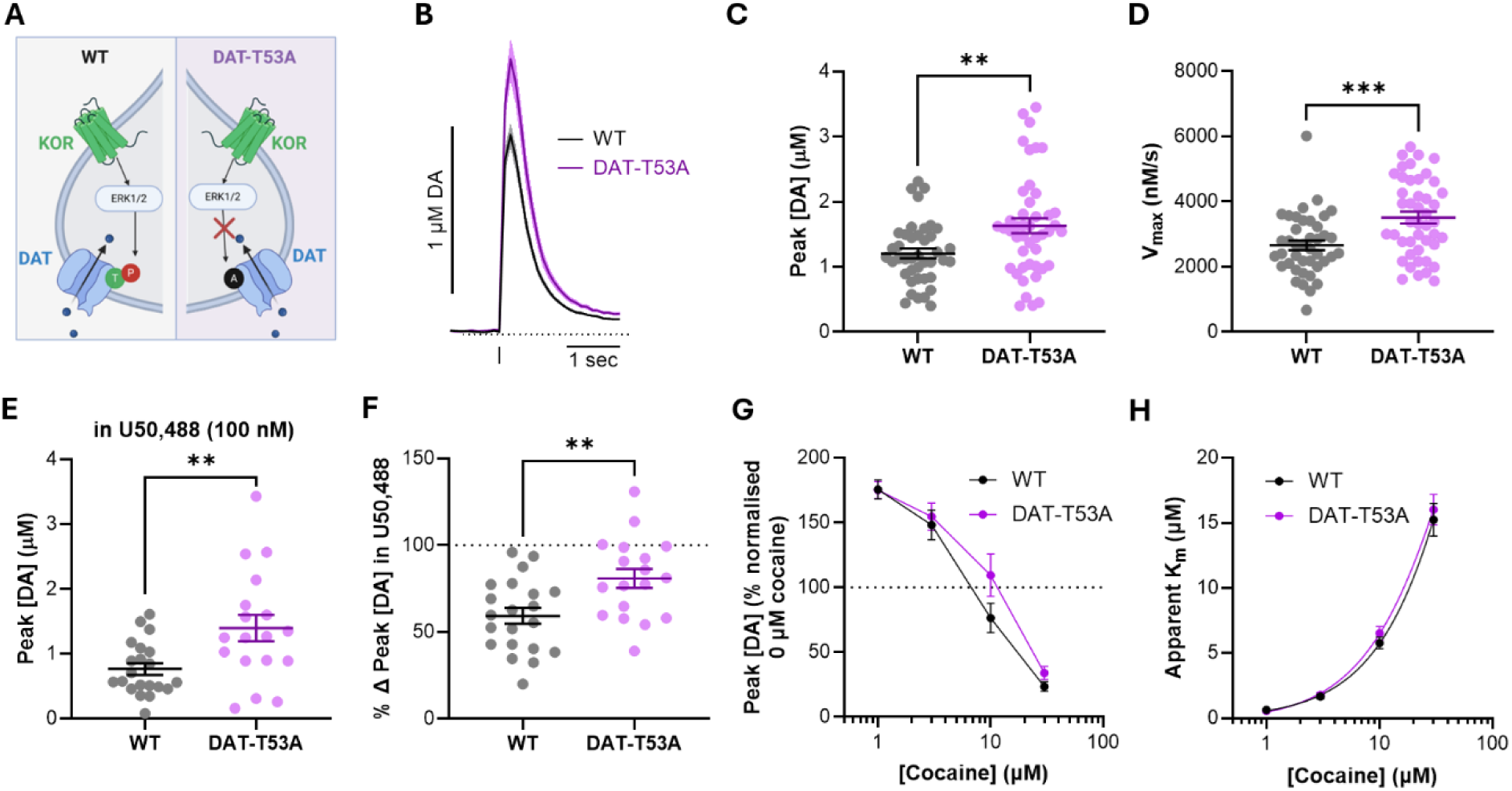
- DAT-T53A mice show altered baseline DA dynamics and blunted KOR responses. **(A)** Schematic describing DAT-T53A mutation. Replacing threonine-53 with an alanine renders this site insensitive to KOR-mediated T53 phosphorylation. **(B)** Average ± SEM WT (black) and DAT-T53A (purple) DA release events, evoked by a single electrical pulse in drug-free conditions. Vertical line indicates electrical stimulation timepoint, horizontal dotted line indicates 0 µM [DA]. **(C)** Average ± SEM peak evoked NAc [DA] in WT and DAT-T53A slices. **(D)** Average ± SEM DA uptake in WT and DAT-T53A NAc, assessed through V_max_. (**E)** Average ± SEM peak [DA] following U50,488 in WT and DAT-T53A slices. **(F)** U50,488-mediated effect on peak [DA], represented as percentage change in peak [DA] ± SEM in WT and DAT-T53A. **(G)** Peak evoked [DA] normalized to peak evoked [DA] at 0 μM cocaine across cocaine concentrations. **(H)** Average apparent K_m_ values across different cocaine concentrations. Data are pooled from drug-free data from Figures 3 and 6. N = 11 WT and 10 DAT-T53A mice. * p ≤ 0.05, ** p ≤ 0.01, *** p ≤ 0.001.

To assess whether there are altered basal DA dynamics in DAT-T53A mice, we compared evoked DA between DAT-T53A and WT animals. We collated drug-free WT and DAT-T53A single-pulse data present throughout **Figures 3 and 6**, and **Figure 5B** represents the resulting average of these events. DAT-T53A animals show higher peak [DA] (**Figure 5B,C**, unpaired t-test: *t*(84) = 3.027, *p* = 0.0033, *n* = 41 WT and 45 DAT-T53A), and faster DA uptake compared to WT controls (**Figure 5B,D**, unpaired t-test: *t*(84) = 3.677, *p* = 0.0004, *n* = 41 WT and 45 DAT-T53A). We further tested whether the effect of U50,488 in suppressing DA release is different between WT and DAT-T53A animals. We found that, in the presence of U50,488, peak evoked [DA] is greater in DAT-T53A animals (**Figure 5E**, unpaired t-test: *t*(37) = 3.001, *p* = 0.0048, *n* = 21 WT and 18 DAT-T53A) and found that U50,488 has a significantly smaller effect in DAT-T53A animals (**Figure 5F**, unpaired t-test: *t*(37) = 3.055, *p* = 0.0042, *n* = 21 WT, 18 DAT-T53A). We detected no significant effect of genotype on cocaine effects on normalized peak [DA] (**Figure 5G**, two-way ANOVA, effect of genotype: *F*(1,180) = 2.972, *p* = 0.0864, *n* = 21 WT, 27 DAT-T53A). Finally, we tested whether the ability of cocaine to inhibit DAT was different between these genotypes. We found no statistically significant effect of genotype when comparing apparent K_m_ between WT and DAT-T53A mice (**Figure 5H**, two-way ANOVA, effect of genotype: *F*(1,184) = 0.6868, *p* = 0.4083, *n* = 21 WT, 27 DAT-T53A). In summary, DAT-T53A mice have increased DA release and uptake, and reduced KOR-mediated suppression of DA release.

**Figure 6.**
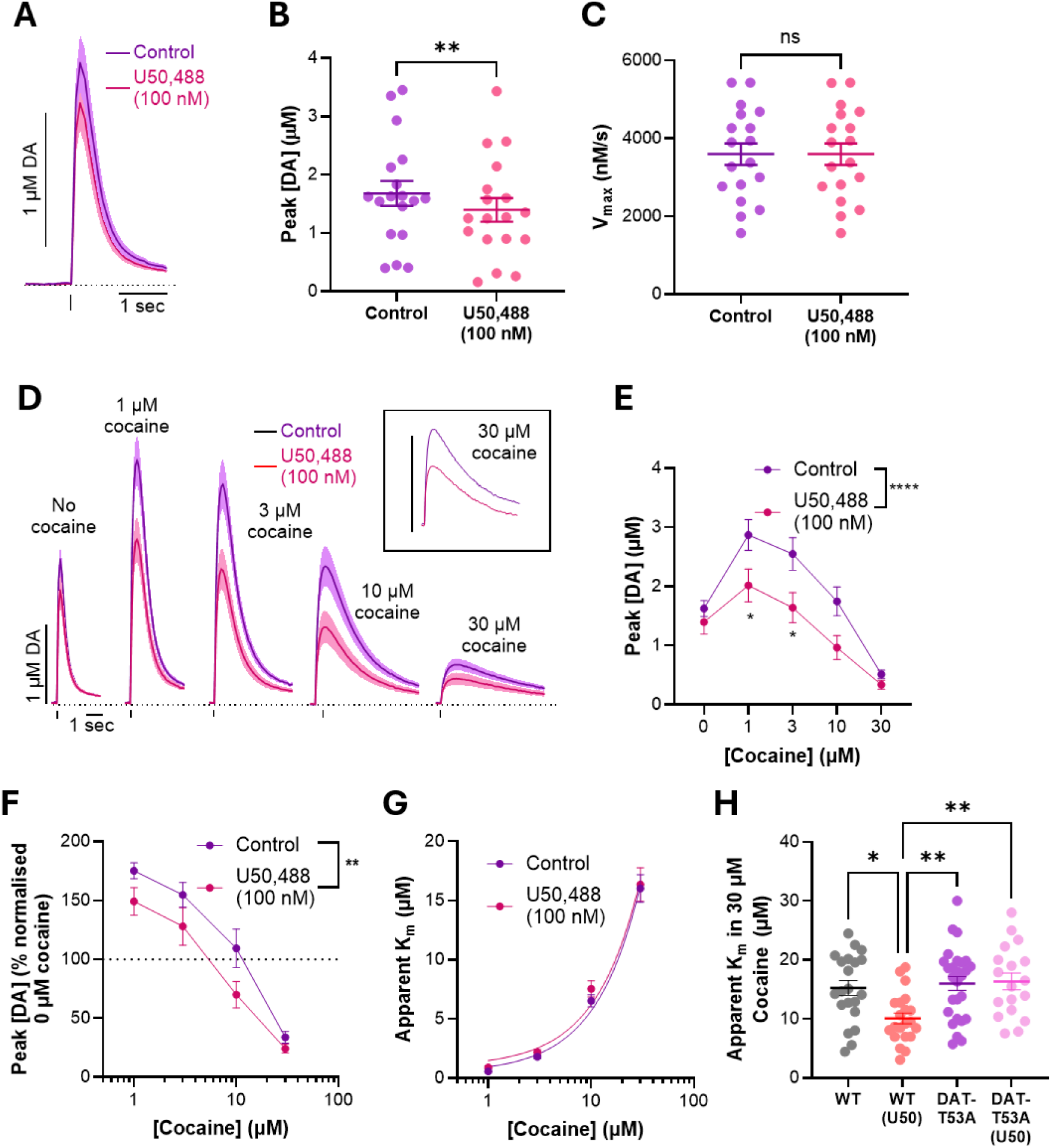
- Phosphorylation of DAT-T53 residue is required for KOR-mediated changes to cocaine uptake effects. **(A)** Average ± SEM DAT-T53A DA release events in drug-free conditions (Control, purple line) and in the presence of the KOR agonist U50,488 (100 nM, pink line). Vertical line indicates electrical stimulation timepoint, horizontal dotted line indicates 0 µM [DA]. **(B)** DAT-T53A peak evoked [DA] in drug-free and U50,488 conditions. **(C)** Maximal velocity of DA uptake (V_max_) in Control and U50,488 conditions. **(D)** Average DAT-T53A DA release events evoked in increasing concentrations of cocaine (0, 1, 3, 10, 30 μM), in the presence or absence of 100 nM U50,488. Inset: magnified representation of average DA release events in 30 μM cocaine. Bar = 0.5 μM [DA]. **(E)** Peak evoked [DA] across cocaine concentrations. **(F)** Peak evoked [DA] normalized to peak evoked [DA] at 0 μM cocaine across cocaine concentrations. **(G)** Average apparent K_m_ values across different cocaine concentrations. **(H)** Average apparent K_m_ values at 30 µM cocaine in WT and DAT-T53A slices, in the presence and absence of U50,488 (U50). n = 26 control, 18 U50,488 slices, N = 5 male, 5 female mice. * p ≤ 0.05, ** p ≤ 0.01.

### Phosphorylation of DAT-T53 residue is required for KOR-mediated DAT tolerance to cocaine

We then tested the effects of the KOR agonist U50,488 (100 nM) on NAc DA release and uptake in the DAT-T53A mice. Similar to WT animals, U50,488 suppressed peak [DA] (**Figure 6A,B**, paired t-test: *t*(17) = 3.060, *p* = 0.0071, *n* = 18), without significantly impacting the rate of DA uptake V_max_ (**Figure 6A,C**, paired t-test: *t*(17) = 0.3681, *p* = 0.7174, *n* = 18).

In slices from DAT-T53A mice, we superfused increasing concentrations of cocaine in the presence or absence of U50,488 (**Figure 6D**), and assessed changes to peak [DA] (**Figure 6E,F**) and apparent K_m_ (**Figure 6G**). As in WT animals, we further compared the effect of U50,488 in the presence of cocaine on peak [DA], and found that U50,488 suppressed peak [DA] (**Figure 6E**, two-way ANOVA, effect of U50,488: *F*(1,210) = 18.06, *p* < 0.0001, n = 26 control, 18 U50,488), with significant post hoc effects at 1 μM and 3 μM cocaine (**Figure 6E**, Šídák’s multiple comparisons test, 1 μM cocaine: *p* = 0.0327, 3 μM cocaine: *p* = 0.0189). We further normalized peak [DA] values across the various cocaine concentrations to the peak [DA] value in the absence of cocaine, and surprisingly found that U50,488 suppressed peak [DA] in the presence of cocaine (**Figure 6F**, two-way ANOVA, effect of U50,488: *F*(1,168) = 10.11, *p* = 0.0018, *n* = 26 control, 18 U50,488). However, when testing the ability of DA to bind to the DAT in the presence of cocaine, U50,488 did not alter apparent K_m_, suggesting phosphorylation of the T53 residue on the DAT is required for KOR-mediated changes to cocaine tolerance (**Figure 6G**, two-way ANOVA, effect of U50,488: *F*(1,168) = 0.9611, *p* = 0.3283, *n* = 26 control, 18 U50,488 slices).

We further tested these effects in male DAT-T53A drug-naïve mice *in vivo* with the KOR agonist U69,593, similar to **Figure 2** (s.c., 0.1 mg/kg**, Supplementary Figure 1A**). In DAT-T53A animals, pretreatment with U69,593 did not significantly suppress peak [DA] (**Supplementary Figure 1B**, unpaired t-test: *t*(32) = 0.8154, *p* = 0.4209, *n* = 16 vehicle and 18 U69,593 slices). Uptake was unchanged following U69,593 pretreatment (**Supplementary Figure 1C**, unpaired t-test: *t*(32) = 0.3893, *p* = 0.6996, *n* = 16 vehicle and 18 U69,593 slices). Finally, the effects of cocaine were not significantly different between DAT-T53A animals pre-treated with vehicle or U69,593 (**Supplementary Figure 1D**, two-way ANOVA, effect of U69,593: *F*(1,50) = 0.1748, *p* = 0.6776, *n* = 6 vehicle and U69,593 slices).

To more directly address the requirement of the T53 residue for KOR-mediated DAT tolerance to cocaine, we compared apparent K_m_ from evoked [DA] transients in the presence of 30 μM cocaine between WT and DAT-T53A animals, in the presence and absence of U50,488 (**Figure 6H**). We show a significant effect of treatment (**Figure 6H**, one-way ANOVA, effect of treatment: *F*(3,83) = 5.831, *p* = 0.0012) and again show significant differences between apparent K_m_ in WT slices in the presence and absence of U50,488, and no significant difference between apparent K_m_ in DAT-T53A slices in the presence and absence of U50,488 (**Figure 6H**, Tukey’s multiple comparisons test, WT vs WT (U50,488): *p* = 0.0186, DAT-T53A vs DAT-T53A (U50,488): *p* = 0.9970). Comparing across genotypes, we show no significant differences in apparent K_m_ between both DAT-T53A conditions with WT in the absence of U50,488 (**Figure 6H**, Tukey’s multiple comparisons test, WT vs DAT-T53A: *p* = 0.9646, WT vs DAT-T53A (U50,488): *p* = 0.9251). Conversely, there are significant differences between WT in the presence of U50,488 and both DAT-T53A conditions (**Figure 6H**, Tukey’s multiple comparisons test, WT (U50,488) vs DAT-T53A: *p* = 0.0025, WT (U50,488) vs DAT-T53A (U50,488): *p* = 0.0041). Put together, these comparisons more directly highlight that phosphorylation of the T53 residue of the DAT is required for DAT tolerance to cocaine.

## Discussion

In summary, we found that inhibiting KORs prevents escalation of cocaine use, and the development of DAT tolerance to cocaine following cocaine self-administration. We further found that an acute application of a KOR agonist is sufficient to reduce the inhibition of DA reuptake by cocaine in NAc core slices. We replicated this finding in NHP NAc core, highlighting the translatability of this finding, and showed its dependence on pT53-DAT in mice.

We have previously demonstrated that in rats, high-dose, binge-like cocaine self-administration procedures led to escalated intake rates across self-administration sessions ^9,11,64,67^ and subsequent reductions in the ability of cocaine to inhibit DA reuptake ^9–11^, suggesting that the escalating rate of intake is driven by a molecular tolerance of the DAT. Alongside these changes, cocaine self-administration increased KOR activity in the NAc core and prodynorphin expression in the VTA and NAc ^18^, and treating these animals with a combination treatment of the DA releaser phenmetrazine and the KOR antagonist nBNI prevented tolerance of the DAT to cocaine ^10^. We now show that pre-treatment with nBNI alone is sufficient to prevent these effects, suggesting that the Dyn/KOR system underlies the tolerance to cocaine. We further demonstrate that higher cocaine intake rates and increased DAT tolerance to cocaine occur simultaneously (**Figure 1**), and that KOR antagonism can lower intake rates while preventing the development of DAT tolerance to cocaine. Consequently, we propose that these findings are consistent with DAT tolerance contributing to increases in cocaine consumption and further suggest that KOR-driven tolerance at the DAT contributes to escalating intake.

Our work more directly reinforces these findings by demonstrating that a single superfusion of a KOR agonist on acute slices is sufficient to cause tolerance of the DAT to cocaine. As we can induce tolerance in acute NAc slices, these effects are localized to the NAc and are likely to be largely mediated by DA terminal KORs interacting with the DAT. A further important consequence of these findings is the timescale of onset of these effects. As mentioned previously, DAT tolerance to cocaine and engagement of the KOR/Dyn system are described as emerging during chronic drug administration. Our findings suggest that KOR’s impact on the DAT can occur with a single application of KOR agonist on acute slices, or a single s.c. injection of a KOR agonist, suggesting these effects are transduced through rapid mechanisms that may not require multiple insults to be engaged. Phosphorylation of T53 has been shown to occur within two hours following systemic U69,593 administration ^56^, sixty seconds following methamphetamine exposure in rat striatal synaptosomes ^68^ or within an hour for cell culture models exposed to methamphetamine ^68^. The persistence of T53 phosphorylation, and its time course in intact terminals, remain uncharacterized and future work should address the temporal dynamics of this process.

We replicated well-documented findings from our lab and others that KOR activation suppresses stimulated DA release in NAc slices ^23,24,26,44,69,70^. We further showed that KOR activation did not change NAc DA uptake in this preparation. This is in line with other published striatal slice work ^69,70^, as well as *in vivo* reports employing microdialysis ^35^ and FSCV ^28^. However, it is important to note that these findings conflict with data from cell expression systems ^32^, striatal synaptosomes ^33,34^, and *in vivo* experiments ^26,34^, which suggest KOR activation increases DA uptake. Consequently, reports of KOR-mediated changes to DA uptake are not consistent throughout the literature; however, our findings are aligned with those from other acute slice experiments. It is worth noting that, while our lab does not see uptake changes in *ex vivo* slices ^23,24,44^, we are able to see increased DA uptake *in vivo* in spontaneous NAc DA signals with the fluorescent DA sensor dLight ^26^. This suggests that differences in methodological approaches may underlie these conflicting findings.

In addition, we tested whether we could replicate these findings in the NAc core of drug-naïve NHPs. Indeed, we showed that KOR activation led to a significant reduction in NAc DA release and decreased the ability of the dual DAT/NET inhibitor nomifensine to inhibit DA uptake. Within the NAc, nomifensine behaves similarly to cocaine, as FSCV studies have shown that chronic cocaine self-administration produces tolerance to both cocaine- and nomifensine-mediated uptake inhibition ^9^. This suggests that these findings are conserved across the species tested and lend translational validity to these mechanisms. This is particularly important given the often-inconsistent findings between rodent and human studies regarding KOR involvement in cocaine use disorder. For example, human post-mortem and imaging studies have either shown ^71,72^, or been unable to find ^73^ changes in KOR binding in individuals with CUD. Furthermore, KOR antagonism has not been shown to be efficacious in reducing subjective cocaine craving or cocaine use in human trials ^74,75^. Better understanding the conserved mechanisms between rodents and humans will allow for the identification of translationally relevant targets.

We employed a mouse line with a substituted alanine for the usual threonine at position 53 (DAT-T53A) to investigate the role of this phosphorylation site on the changes in cocaine effects. We showed that the DAT-T53A mutation abolishes KOR-induced changes in cocaine potency using *ex vivo* FSCV, highlighting the requirement for the T53 site to mediate these effects. However, the mechanism behind pT53 altering the ability of cocaine to inhibit uptake is not understood. Several residues in transmembrane domains 2 and 3 of mouse DAT have been linked to the potency of cocaine inhibition ^76,77^ and DAT phosphorylation has been linked to many outcomes (reviewed in ^53^), including modifying uptake, trafficking, and DAT degradation. One molecular mechanism that has been linked to DAT tolerance to cocaine is the formation of DAT oligomer complexes ^78^; however, it is not known whether KOR activation or pT53 regulates DAT oligomerization. Future studies should attempt to narrow down the mechanism underlying this effect.

DAT-T53A mice were previously shown to be insensitive to KOR-mediated locomotor suppression and conditioned place aversion ^56^. Interestingly, we showed a reduction in the effect of a KOR agonist on NAc core DA release compared to WT, highlighting a similar insensitivity to KOR activation at the terminal level. While this finding matches the previously published behavioral data, the mechanism of this change is unknown. The DAT mediates a depolarizing conductance during DA uptake that may influence terminal excitability and thereby oppose the inhibitory effects of KOR activation ^79,80^. Additionally, KORs mediate several intracellular G-protein dependent and independent signaling cascades ^81,82^. It is possible that the basal changes to release and uptake in DAT-T53A animals may shift intracellular signaling molecules that have a knock-on effect on KOR-mediated effects on DA release. Indeed, D_3_ autoreceptor inhibition and activation of D_2_ autoreceptors have been shown to enhance or inhibit cocaine potency at the DAT, respectively ^83^. Future work should test other GPCR signaling systems present in DA terminals to assess whether other G_i/o_ receptor systems (D_2_-like or GABA_B_) are similarly affected in DAT-T53A animals.

We further observed that U50,488 mediates a small suppression of evoked [DA] in DAT-T53A mice (**Figure 6B**). Interestingly, in these animals, U50,488 mediates a significant suppression of peak evoked [DA] in the presence of cocaine (**Figure 6F**). This contrasts with WT animals which present with a larger suppression of DA release mediated by U50,488 (**Figure 3A,B**) but no further effect of KOR activation in the presence of cocaine (**Figure 3F**). We hypothesize that this surprising finding may again be evidence of an interaction between DAT-mediated depolarizing and KOR-mediated inhibitory conductances. A faster-cycling T53A DAT could deliver more depolarizing charge to DA terminals, functionally opposing inhibition from KORs. In the presence of cocaine, the DAT’s opposing conductance is blocked, allowing for the emergence of KOR-mediated inhibition of peak [DA]. In WT mice the DAT is less active, resulting in a smaller depolarization and less opposition against KOR-mediated inhibition.

KOR agonists have been shown to increase cocaine conditioned place preference (CPP) in WT animals, but not in DAT-T53A mice ^56^. Indeed, pT53-DAT has been shown in several studies to increase the affinity of cocaine for the DAT and modulate cocaine-associated behaviors ^56,57,84–86^. Our results may initially seem contradictory, as we show KOR agonism reduces cocaine’s ability to inhibit DA uptake in WT animals, suggesting that the animal may experience a reduced DA elevation in response to cocaine. This may lead to the conclusion that, with KOR activation, cocaine is experienced as less rewarding, with an expectation that CPP be reduced with KOR agonism. However, as KOR activation is aversive, it may be possible that the subjective experience of receiving an aversive stimulus followed by cocaine, regardless of the “potency” of the experienced cocaine administration, could be more salient and rewarding than a dose given in the absence of aversion ^29,87,88^. This may explain why DAT-T53A animals do not show enhanced CPP following KOR agonist administration. As there is a reduced efficacy of KOR agonists and a lack of conditioned place aversion, it is possible DAT-T53A animals experience little aversion to KOR agonism. This suggests that, in DAT-T53A animals, the experienced valence of cocaine dose is unaffected by a KOR-mediated aversive state.

Previous findings in DAT-T53A mice reported hyperlocomotion and enhanced exploratory behaviors, suggesting elevated DA release ^57^. Consistent with this interpretation, we confirmed an increase in electrically-stimulated DA release in DAT-T53A mutants. Additionally, we have shown that DA uptake is elevated in DAT-T53A animals when compared to WT. In support of this, one study has shown increased DA uptake in LLC-PK_1_ cells expressing DAT-T53A assessed through rotating-disk electrode voltammetry ^68^. However, these findings contrast with previous studies using synaptosomes obtained from DAT-T53A, which do not find a difference in DAT-mediated uptake in the DAT-T53A mutants ^56,57^. We predict this mismatch may be due to methodological differences, as acute slice measurements rely on endogenous DA and retained terminal architecture. We hypothesize that the accelerated transport kinetics of the T53A DAT leads to enhanced vesicular refilling and recycling, enabling terminals to sustain greater electrically-stimulated release. Consistent with this idea, DAT-T53A mutants exhibit higher evoked absolute DA concentrations during cocaine exposure (**Figures 3E and 6E**), which may indicate that the terminal can accommodate elevated release demands, even while DA uptake is blocked by increasing cocaine concentrations. It is worth noting that cocaine concentration-response curves show increases followed by a decline at higher concentrations. A balance of many mechanisms is likely to cause these effects, principally DA reuptake inhibition, cocaine enhancing a synapsin-dependent release mechanism (though this has been recently challenged ^89^), and autoreceptor activation ^70,90–92^.

A major limitation of our study is the exclusive use of male animals for some datasets, particularly **Figure 1**. Female rodents have been shown to be less sensitive to KOR agonist effects on antinociception, motivated behavior, and depressive-like behavior ^93–96^. DA release *in vivo* ^94^ and in striatal tissue ^97^ shows smaller reductions in female animals with KOR agonists; however, *ex vivo* work in NHP ^24,58^ and rodents ^39^ show no sex differences. We note that the keystone experiments exploring KOR-mediated DAT tolerance to cocaine (**Figures 3, 5, 6**) were performed in male and female mice with no significant sex effects seen throughout; consequently, these KOR-DAT interactions are likely to be present in both male and female mice. However, as most behavioral data suggests that females have a reduced effect of KOR agonists, how this mechanism differentially impacts behavior in male and female animals remains a valid and unanswered question we aim to address in the future. We further note that the only sex effect we identified in our data was a significant interaction between sex and nomifensine on apparent K_m_ changes in non-human primate slices in **Figure 4K**. While significant, we caution overinterpreting this finding as it results from 3 male and 4 female data points per condition and was not powered to detect sex differences.

In conclusion, we demonstrated that inhibiting KORs prevents escalation of cocaine use, and the development of DAT tolerance to cocaine following cocaine self-administration. NAc core KORs mediate cocaine-mediated inhibition of DA uptake and this process requires phosphorylation of the T53 residue, narrowing the Dyn/KOR-DAT signaling axis to a discrete target and suggesting a molecular pathway that underlies cocaine tolerance. We further show that this effect is present in NHPs, highlighting that these pathways are conserved from rodents to primates and are putative targets for development of treatments for substance-use disorders.

## Data availability

Data supporting the findings of this study are available from the corresponding author upon reasonable request.

## Author contributions

E.F.L., P.M.E., K.M.H., and S.R.J. conceptualized and designed the study. E.F.L., P.M.E., A.M.C., K.R.B., M.H.D., J.H.S., and K.M.H. performed experiments and analyzed data. K.A.G. and V.C.C.C. provided non-human primate tissue and associated resources. L.D.J. and S.R. generated and characterized the DAT-T53A mouse line and contributed to interpretation. E.F.L. and S.R.J. wrote the manuscript. All authors reviewed, edited, and approved the final manuscript.

---

**Supplementary Figure 1.**
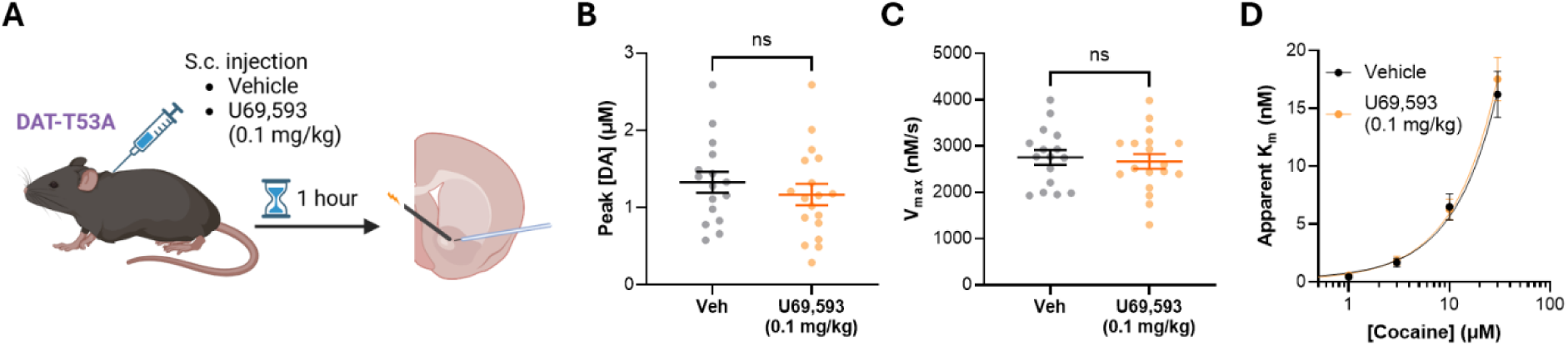
- *In vivo* pretreatment with KOR agonist does not induce DAT tolerance in DAT-T53A mice. **(A)** Schematic of experimental set-up. Male DAT-T53A mice were injected with the KOR agonist U69,593 (0.1 mg/kg) or vehicle (20% propylene glycol in saline). Following a 1 hour period, animals were sacrificed and acute NAc core striatal slices were obtained for FSCV. **(B)** Average ± SEM peak evoked [DA] in vehicle (black) and U69,593 (yellow) conditions. **(C)** Maximal velocity of DA uptake (V_max_) in vehicle and U69,593 conditions. **(D)** Average apparent K_m_ across cocaine concentrations in vehicle and U69,593 treated animals. n = 6-16 vehicle, 6-18 U69,593 slices. N = 6 vehicle, 6 U69,593-treated mice.

